# Characterization of mouse *Bmp5* regulatory injury element in zebrafish wound models

**DOI:** 10.1101/2021.05.17.444306

**Authors:** Ian S. Heller, Catherine A. Guenther, Ana M. Meireles, William S. Talbot, David M. Kingsley

## Abstract

Many key signaling molecules used to build tissues during embryonic development are re-activated at injury sites to stimulate tissue regeneration and repair. Bone morphogenetic proteins provide a classic example, but the mechanisms that lead to reactivation of BMPs following injury are still unknown. Previous studies have mapped a large “*injury response element*” (*IRE*) in the mouse *Bmp5* gene that drives gene expression following bone fractures and other types of injury. Here we show that the large mouse *IRE* region is also activated in both zebrafish tail resection and mechanosensory hair cell injury models. Using the ability to test multiple constructs and image temporal and spatial dynamics following injury responses, we have narrowed the original size of the mouse *IRE* region by over 100 fold and identified a small 142 bp minimal enhancer that is rapidly induced in both mesenchymal and epithelial tissues after injury. These studies identify a small sequence that responds to evolutionarily conserved local signals in wounded tissues and suggest candidate pathways that contribute to BMP reactivation after injury.

## 1 Introduction

Regenerating skeletal tissue after injuries is essential to millions of people who fracture a bone each year (Tosounidis *et al*. 2009, Tsang *et al*. 2009, Farr *et al*. 2017). For many fractures, stabilizing the broken bone allows for repair by an innate process of localized bone growth. However, approximately 1-8% of bone fractures do not heal (Mills *et al*. 2017, Ekegren *et al*. 2018). Better methods of stimulating repair at fracture sites would be of great use for such injuries. Synthetic growth factors such as Bone Morphogenetic Proteins (BMPs) are able to stimulate bone growth and are widely used in orthopedic applications (Axelrad & Einhorn 2009). However, because BMPs are potent signaling molecules with multiple effects on proliferation, differentiation and development, exogenous BMPs must be applied carefully to avoid undesired complications including heterotopic bone growth, inflammation, and other unwanted effects (Ronga *et al*. 2013, Barcak & Beebe 2017).

Expression of the body’s own BMP molecules is induced specifically at sites of bone fractures, and many of the steps in fracture healing resemble the sequence of events that occur during embryonic bone formation (Einhorn 1998, Goldring *et al*. 2006). Tissue regeneration in many organisms is thought to take place by reactivation of the same signaling molecules used to control tissue formation during embryonic development (Darnet *et al*. 2019, Yang & Kang 2019). However, the mechanisms that lead to reactivation of key signaling molecules at an injury site are still poorly understood. Adult signal reactivation may occur either through the same DNA enhancers that control embryonic expression of a key signaling gene, or through separate, injury-specific regulatory enhancers (Rodriguez & Kang 2019, Suzuki & Ochi 2020). A better understanding of the mechanisms that regulate local BMP expression following injury may suggest new ways to improve localized, controlled, and clinically useful BMP expression for adult tissue regeneration.

Our understanding of embryonic and adult BMP regulation in mammals is currently most advanced for the *Bmp5* gene. A large collection of coding and regulatory mutations at the mouse *Bmp5* locus were isolated during studies of the classic morphological trait, “short-ear” (Lynch 1921, Russell 1951, Rinchik *et al*. 1985, Kingsley *et al*. 1992, Marker *et al*. 1997). Mice homozygous for null mutations at the *Bmp5* (*short ear)* locus are viable and fertile, but show changes in the size, shape, or number of many specific skeletal structures. The characteristic morphological changes in the mice have been traced to localized changes in particular skeletal condensations, cartilage, and bone elements during embryonic and early postnatal development (E. Green & McNutt 1941, E. Green & Green 1942). In addition to these programmed changes in specific mouse skeletal structures, *Bmp5/short ear* mice also show reduced formation of a regenerative fracture callus following rib fracture in adulthood (M. C. Green 1958). The combination of both anatomical changes and fracture healing defects in *short ear* mice provided the first genetic evidence that BMPs are required both for normal skeletal development and for regeneration of skeletal structures after injury (Kingsley *et al*. 1992, Kingsley 1994).

Regulatory mutations at the *short ear* locus suggest that key developmental enhancers of the gene are distributed over hundreds of kilobases of DNA found upstream, intronic, and downstream of *Bmp5* coding exons (DiLeone *et al*. 1998). Large-scale *Bmp5* enhancer scans in transgenic mice have identified dozens of developmental enhancers that control the expression of *Bmp5* in particular embryonic skeletal structures or soft tissues (DiLeone *et al*. 1998, 2000, Guenther *et al*. 2008, 2015). Similar studies at postnatal stages have also identified a 17.8 kb “*injury response element*” (*IRE*) that drives gene expression following adult rib fractures (Guenther *et al*. 2015). The *IRE* that responds to unpredictable rib injuries is distinct from the developmental enhancers that drive stereotyped *Bmp5* expression during normal rib formation. Thus, while development and repair activate the same gene, the control sequences for developmental and injury-induced expression in the same tissue are different. Interestingly, the same *IRE* region that drives expression following adult rib fracture also drives expression following multiple other types of adult injury, including fracture of tibia, naphthalene-induced injury of lung epithelium, or mechanical injuries of skin epithelium (Guenther *et al*. 2015).

To address whether the *IRE* contains a single functional element that drives injury response in both mesenchymal and epithelial tissues, or might contain clustered but distinct enhancers for injury in multiple tissues (Kang *et al*. 2016), we sought to further narrow the large 17.8 kb *IRE* region. Our previous approach of serial transgenic assays in mice was slow, labor-intensive, and not amenable to live-imaging during injury responses. In contrast, transgenic zebrafish are easy to construct, offer large clutches, transparent tissues, and well-established wound models. Here we test the ability of the *IRE* region to activate gene expression following injury in zebrafish and characterize the dynamics and minimal sequences required for *IRE* expression.

## 2 Material and Methods

### 2.1 Fish

Wildtype (TL, AB/TU) and transgenic male and female zebrafish were maintained at 28°C with a 14hr:10hr day:night light cycle and fed twice daily. Fish were anesthetized before surgery, experimental treatment, and during imaging with 0.016% (w/v) tricaine methanesulfonate. The transgenic lines *tg(ΔNp63:Gal4)* (Rasmussen *et al*. 2015), *tg(UAS-E1B:NTR-mCherry)* (Goll *et al*. 2009) (ZIRC Catalog ID: ZL1473), *tg(mpeg1:Gal4)* (Ellett *et al*. 2011), and *tg*(*NBT:dsRed*) (Peri & Nüsslein-Volhard 2008) have been described previously. The neutrophil specific expression vector *pTol2 lyz:mCherryCAAX:pA;cmcl2:GFP* vector was assembled using multisite Gateway (Kwan *et al*. 2007) and the following vectors: p5E-*lyzC*, pME-mCherryCAAX, p3E-polyA (Shiau *et al*. 2013) and pDestTol2CG2 (Kwan *et al*. 2007). Tg*(lyz:mcherryCAAXpA;cmcl2:GFP)* fish were generated by co-injecting 1-cell stage wildtype embryos with 12-25 pg of neutrophil specific expression vector and 50-100 ng of *Tol2* transposase mRNA. Injected animals with strong green heart signal (from *cmcl2*:*GFP*) and red neutrophils were raised to adulthood and outcrossed to wildtype TL fish to generate a stable line. Fish procedures were done in accordance with protocols approved by the Stanford University Institutional Animal Care and Use Committee.

### 2.2 Enhancer-reporter plasmids

Candidate enhancer sequences were tested in the p5’*Tol2:e1b:eGFP:Tol2* expression vector (Q. Li *et al*. 2010) which contains a minimal *E1B* promoter sequence (Wu *et al*. 1987), the *eGFP* reporter cassette (Cormack *et al*. 1996), and Tol2 flanking repeats to facilitate transgenesis. The vector was modified with the *cmlc2:mCherry* fluorescent heart expression cassette (C.-J. Huang *et al*. 2003) cloned into its *AatII* site to provide an independent marker of transgenesis. Two *Not1* sites were removed from the vector by serial partial *Not1* digestions, Klenow end blunting, and ligation. Then, a new *Not1* site was added upstream of the minimal promoter using a *XhoI/BglII* digestion to linearize the plasmid, followed by ligation with overlapping oligos containing the new *Not1* site (Supplemental Table S1). The 17.8 kb *Not1* restriction fragment containing the mouse *IRE* region from Ph7-lacZ (DiLeone *et al*. 1998) was inserted into the new *Not1* site in the expression vector. Candidate enhancer constructs, *R1, R2, R3* were subcloned from intermediate plasmids built with inserts amplified from subregions of BAC B (Guenther *et al*. 2015) containing the mouse *Bmp5 IRE* region using the primers noted in Table S1.

The regions of sequence conservation contained within construct *IRE R3* and used to build *CS All* were amplified from plasmid pKG301, an existing plasmid containing all the conserved regions within the *IRE*. To build pKG301, 8 sequences (pKG301_rI through pKG301_rVIII) conserved between mouse and human were identified within the *IRE* (Ph7; Ralph J. DiLeone, Russell, and Kingsley 1998) using VISTA (Mayor *et al*. 2000). Each individual conserved region was amplified using primers in Table S1. Three pairs of conserved sequences (pKG301_rII + pKG301_rIII; pKG301_rIV + pKG301_rV; pKG301_rVII + pKG301_rVIII) were stitched together via PCR using primers noted in Table S1. A final round of PCR was used to stitch together pKG301_rI to pKG301_rII+rIII and pKG301_rVI to pKG301_rVII+VIII using primers noted in table S1. PCR product pKG301_r1+rII+rIII was gel purified, digested with *SpeI* and *XbaI*, and cloned into the *XbaI* site of pBS-KS+ to form pKG289. PCR product pKG301_IV+V was gel purified, digested with *SpeI* and *XbaI*, and cloned into the *XbaI* site of pKG289 to form pKG292. PCR product pKG301_VI+VII+VIII was gel purified, digested with *EaeI* and *NotI*, and cloned into pKG292 that had been partially digested with *NotI* to form pKG293. The concatenated rI-rVIII insert was liberated from pKG293 using *NotI* and cloned into the *NotI* site of 5’Not1hsplacZ (DiLeone *et al*. 1998) to make pKG301_rI-rVIII. The conserved sequences within *IRE* subclone *R3* were amplified with Gibson primers described in Table S1 and cloned into the zebrafish expression vector.

To build constructs *CSE, IRE(483bp)*, and *IRE(142bp)*, the regions were first amplified from BAC B using primers noted in Table S1, and Gibson cloned into the expression vector. The Gibson primers used on the 3’ side of the inserts included a *Not1* site. To build the insert as a tandem repeat, the insert was reamplified using primers (Table S1) with *Not1* sites included on both sides, followed by restriction cloning the PCR product back into the *Not1* site maintained during the first Gibson cloning reaction. Constructs *IRE(142bp [ATT > CCG]), IRE(130bp [Δ12bp])*, and *IRE(92bp [Δ50bp])* were constructed as in *IRE(142bp)*, except using annealed oligos, noted in Table S1 as the PCR template. The sizes and coordinates of all tested *Bmp5 IRE* regions are shown in Supplemental Table S2.

### 2.3 Transcription factor binding site prediction

Candidate transcription factor binding sites (TFBS) were determined by using tfbsPredict to search the UNIPROBE database (Newburger & Bulyk 2009) of position weight matrices (Wenger *et al*. 2013). At the 900 information content threshold the *IRE(142bp)* region contained 6 stretches of DNA with predicted TFBS. A list of potential upstream regulators was intersected with expression data for transcription factors known to be expressed in zebrafish fins after injury, based on *in situ* data collected from the literature and maintained by the Zebrafish Information Network (http://zfin.org/ [2019]). One of the six regions contained a homeobox binding site. This site was predicted to be bound by up to 89 different factors, some of which are known to be expressed after injury (Supplemental Table S3). The engineered mutations of *IRE(142bp [ATT > CCG])* and *IRE(130bp [Δ12bp])* were run through UNIPROBE, using the less stringent “search for TF Binding Sites” method (Newburger & Bulyk 2009) at the recommended 0.45 enrichment threshold and selected based on their ability to disrupt predicted transcription factor binding (Supplemental Table S4).

### 2.4 Tail injury assays and imaging

Transgenic fish were generated by microinjection of test constructs together with 50-100 pg of *Tol2* transposase mRNA into fertilized 1-cell-stage embryos (K. Kawakami 2005). Three day-post-fertilization larvae were screened for *mCherry* fluorescence in the majority of heart cells. Larvae were anesthetized and larval fin folds were amputated at a position immediately caudal to the end of the notochord using a 22.5 degree microknife (FST) and PDMS cutting surface. Fish were then embedded in 1.5% low-melt agarose and imaged on Zeiss LSM 7 microscope with the Plan-Neofluar 10x (numerical aperture 0.30) objective. Time-lapse images were captured at a frequency of 1 z-stack every 10, 7, or 5 minutes for up to 12 hours. Adult fins were amputated using a razor blade approximately half-way along the length of the fin and maintained at 28°C. Adult fish were imaged on a Nikon SMZ1500 outfitted with the Nightsea Stereomicroscope Fluorescence Adapter system and imaged with a Nikon D80 camera.

### 2.5 Neomycin treatment

Neomycin sulfate (Sigma) was dissolved in zebrafish embryo water to a stock concentration of 6 mM and then diluted to a working concentration of 50 µM (Harris *et al*. 2003). Transgenic embryos were generated and screened for heart expression as above, then 5-day post fertilization larvae were incubated in neomycin for 1 hour at 29°C, rinsed in embryo water, and allowed to recover for approximately 0.5 hours before imaging on a Zeiss LSM 7 microscope with the Plan-Apochromat 20x (numerical aperture 0.8) objective.

### 2.6 Morpholino injection

Morpholinos were obtained from Gene Tools (Philomath, OR, USA) and prepared in water as a 1.5 mM stock solution. The sequence of the *spi1b* morpholino antisense oligonucleotide is GATATACTGATACTCCATTGGTGGT (Rhodes *et al*. 2005). For embryo microinjection, morpholinos were diluted in water containing 0.5% phenol red and injected into the embryos of a cross between tg(*IRE R3:GFP*; *cmlc2:mCherry)* and *tg(lyz:mCherry:cmlc2:GFP)* to a final mass of 2-4 ng per embryo. Fish were dechorionated at 2 dpf and sorted for both transgenes using heart reporter expression. Fins were injured as described above, then imaged 24 hours afterward. Images were cropped to a 300 × 600 pixel rectangle centered on the fish and with one edge abutting the wound. In ImageJ, a radius=3 Median filter, then a sigma=2 Gaussian Blur filter were applied to the red channel. Neutrophils were segmented with either Renyi Entropy or Otsu thresholds, then counted with the Analyze Particles tool. Neutrophil counts were checked manually to resolve cases where the two thresholding methods disagreed. To measure the *IRE* response, the green channel threshold was 6000 intensity units, and the area of pixels above that threshold was recorded.

### 2.7 Bat sequencing

Genomic DNA was extracted from a macrobat cell line (*Pteropus alecto*, Sarah Gonzolaes-van Horn) and a microbat tissue sample (*Artibeus jamaicenis*, Lubee Bat Conservancy, Gainesville, FL, USA). Primers flanking the *IRE(142bp)* were designed based on *Desmodus rotundas* sequence. PCR was performed using Phusion polymerase (ThermoFisher) with primers shown in Table S1, standard Phusion conditions, an annealing temperature of 55°C, and 1 minute extension time. The amplified products were gel extracted and cloned using the PCR Blunt II TOPO vector (ThermoFisher). DNA was then sequenced using primers corresponding to the forward and reverse M13 site.

## 3 Results

### 3.1 Mouse injury response element is functional in adult zebrafish

To determine whether zebrafish would be useful to further characterize the mouse *Bmp5* injury response element (*IRE*) we cloned the previously reported *IRE* (*IRE17.8)* region into a *GFP* reporter construct (Fig. 1A), injected the construct into fertilized eggs, and raised transgenic *tg(IRE17.8:GFP)* zebrafish to adulthood. Mosaic founder fish were screened for injury response, and some were bred to transmit the transgene and establish stable transgenic lines. Injury response was tested by resecting caudal fins by clipping the fin at approximately halfway between the hypural bone and the tail end. Repeat imaging of individual fish showed that *IRE17.8* drove robust expression after injury to the caudal fin (Fig. 1B, stable line, Sup. Fig. 1, mosaic founder fish). GFP expression was observed in both the regenerative blastema, as well as the reforming ray and the inter-ray tissue. Fin hemirays that were incidentally broken during manipulation also showed GFP expression surrounding the break (Sup. Fig. 1, arrows).

**Figure 1.**
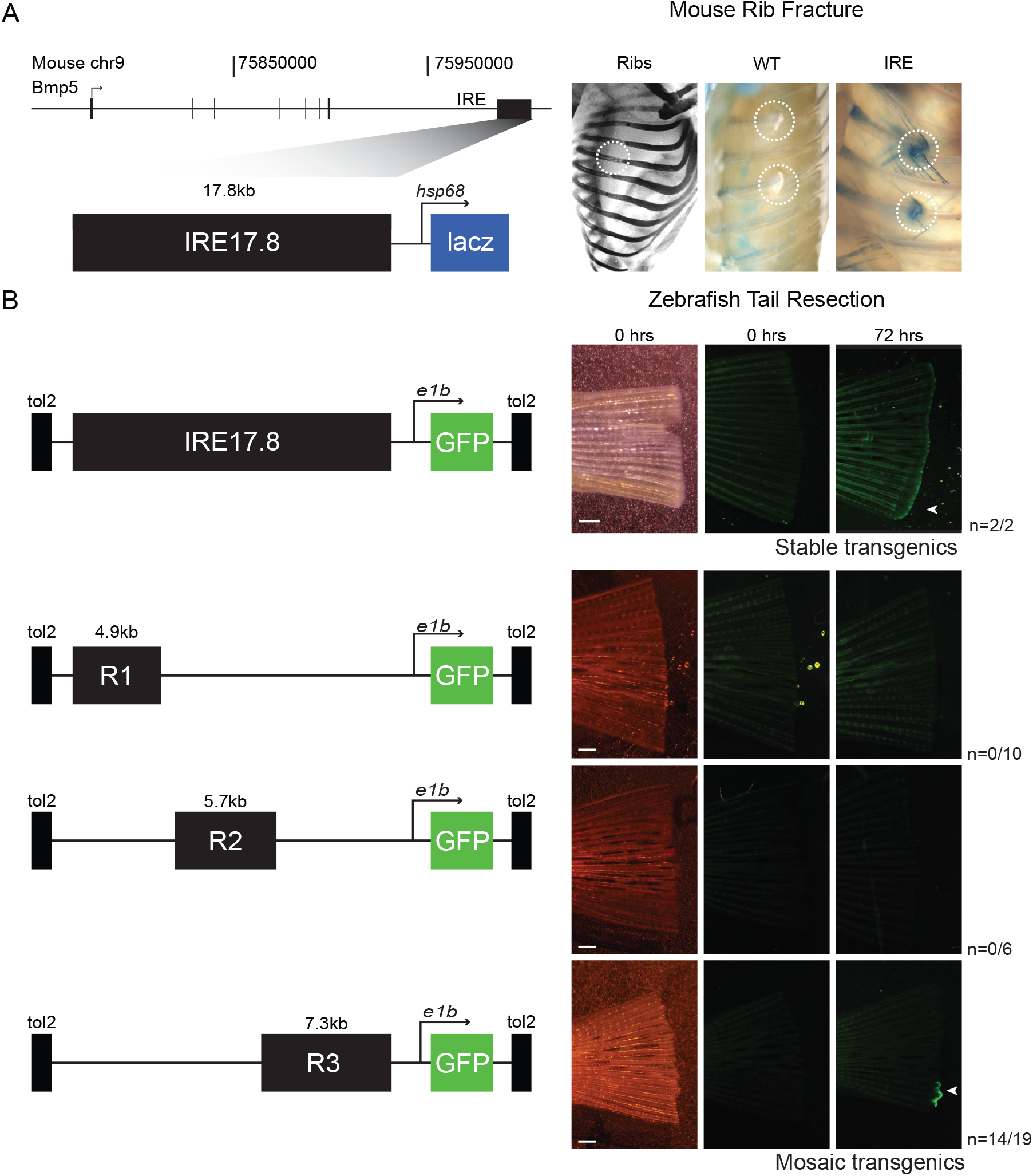
Mouse *Bmp5* injury regulatory region is functional in adult zebrafish caudal fin injury. (A) Schematic of the *Bmp5* locus in mouse genome (mm10 chr9: 75771537-76011386) showing the location of a previously identified injury regulatory region. When cloned upstream of a *LacZ* reporter gene and used to make transgenic mice, this *IRE* region drives expression at the site of adult rib fractures (dotted circles indicating rib breakage in WT and *IRE* mice, modified from Guenther *et al*. 2015). (B) Schematic of reporter constructs used to test injury element in adult transgenic zebrafish (left). Representative caudal fins showing the location of injury and *GFP* expression at 72 hours-post-injury. GFP reporter expression is present at the wound edge of stably transgenic *tg(IRE17.8:GFP)* fish carrying the original *IRE*. When three *IRE* subregions are tested in mosaic transgenic founders, reporter expression is also observed at wound edge of fins with *tg(IRE R3:GFP)* (white arrow), but not *tg(R1:GFP)* or *tg(R2:GFP)*. n = *GFP* positive fish/ total stable (*IRE17.8*) or mosaic (*R1, R2, R3*) transgenic fish screened. Scale bars = 5 mm.

Although typical vertebrates enhancers are less than a few kilobases in length (Yáñez-Cuna *et al*. 2013, L. Li & Wunderlich 2017), the original mouse *IRE* region was a 17.8 kb region (Guenther *et al*. 2015). To further refine the *IRE*, we tested three subclones for their ability to drive expression in the zebrafish caudal fin injury assay. Adult mosaic transgenic *tg(R1:GFP)* and *tg(R2:GFP)* fish did not exhibit significant GFP expression in the wounded tail (Fig. 2B, Sup. Fig. 2). In contrast, *tg(IRE R3:GFP)* fish showed robust expression in fins following injury, in a similar, albeit mosaic, pattern to *tg(IRE17.8:GFP)* fish. Injury-induced expression was confirmed in 2 independent *tg(IRE R3:GFP)* stable lines. These results show that *IRE* is activated in both mouse and zebrafish following tissue injury, and that zebrafish can be used to narrow the *IRE*.

**Figure 2.**
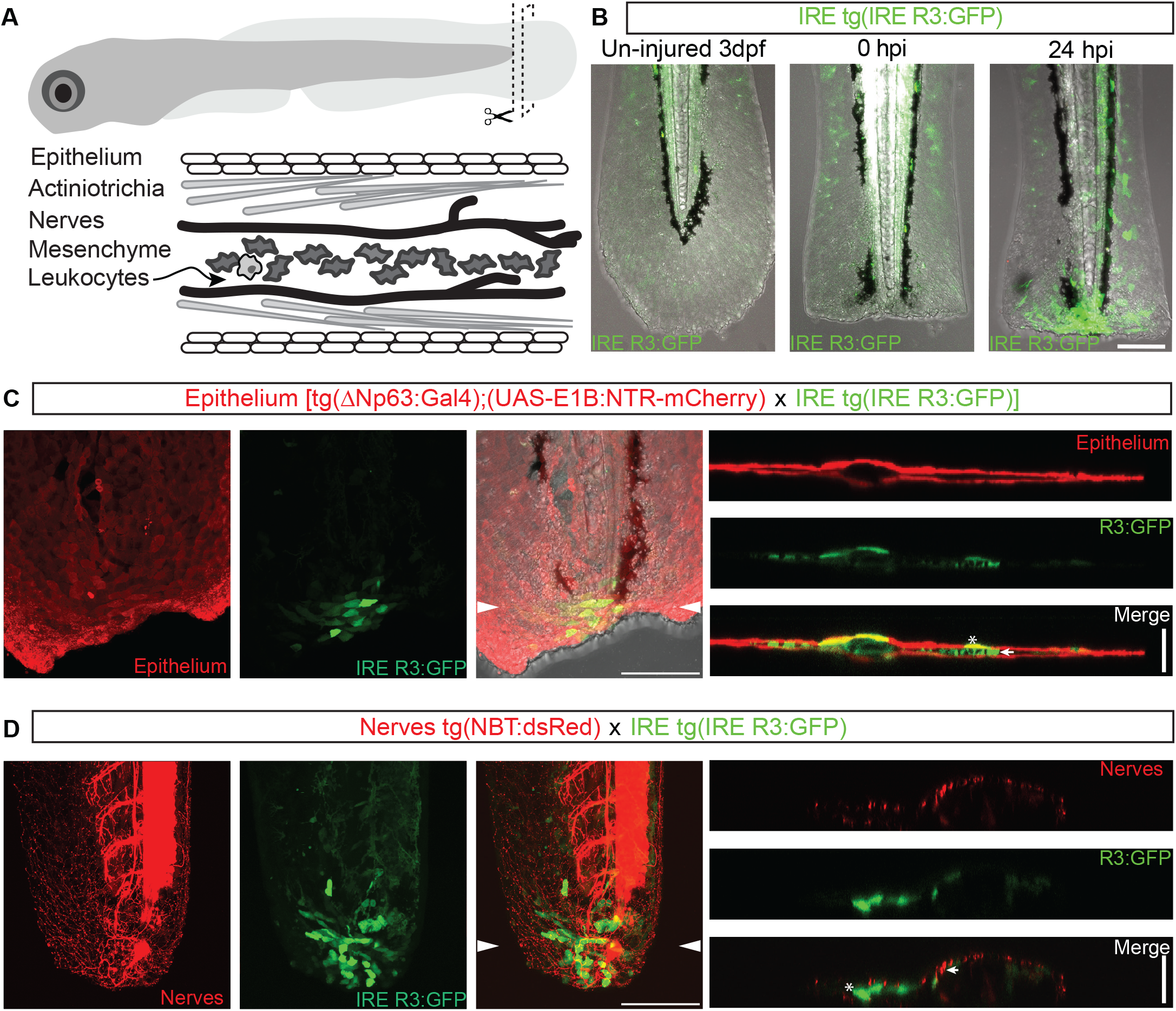
*Bmp5* injury response element reporter expression in stable larval fin fold cell populations. (A) Schematic of larval injury model and the composition of the caudal fin fold. (B-E) Maximum intensity projection confocal images of larvae fin injuries. (B) GFP reporter expression after injury in stable *tg(IRE R3:GFP)* 3 day-post-fertilization larvae. (C) Triple transgenic reporter line *tg(ΔNp63:Gal4; UAS-E1B:NTR-mCherry; IRE R3:GFP)* with epithelial cells marked in red and *IRE R3-GFP* expression shown in green at 24 hpi. Right hand panels show reconstructed cross section of confocal image, at location indicated by white arrowheads. Note that *IRE R3*-driven-*GFP* expression is evident in both epithelial cells, (yellow cells and white asterisk), as well as in mesenchymal cells (green-only interior cells, white arrow). (D) Double transgenic reporter line *tg(NBT:dsRed; IRE R3:GFP)* with neurons marked in red at 24 hpi. Right hand panels show reconstructed cross section of confocal image, at location marked with white arrowheads. Note absence of colocalization between neurons (arrow) and *tg(IRE R3:GFP*) responsive cells (asterisk). Scale bars: lateral views =100 µm, cross sections = 50 µm.

### 3.2 *IRE* activity in larval tissue

Injuries to the fin fold of larval zebrafish also provide a well-studied regeneration model (A. Kawakami *et al*. 2004, Mateus *et al*. 2012). Since larval wounding recapitulates many of the gene expression patterns observed in adult injury (Mathew *et al*. 2009, Yoshinari *et al*. 2009), we next asked whether *IRE R3:GFP* reporter activity could be detected in 3 day post fertilization (dpf) larval zebrafish. The caudal fin fold was clipped at the tip of the notochord of a stable line of larval *tg(IRE R3:GFP)* fish and allowed to recover for 24 hours. The *tg(IRE R3:GFP)* fish exhibited a strong injury response in cells immediately proximal to the wound margin (Fig. 2B-D). Additionally, in some wounds, *GFP* expression was variably detected in more distal cells. Live imaging of *tg(IRE R3:GFP)* fish shows *GFP* detectable above background between 3 and 7.5 hours-post-injury (hpi), with cells adjacent to the wound expressing *GFP* before more distant cells (mean: 3.3 hours, n=4) (Fig. 3A-B, Movie 1-2).

**Figure 3:**
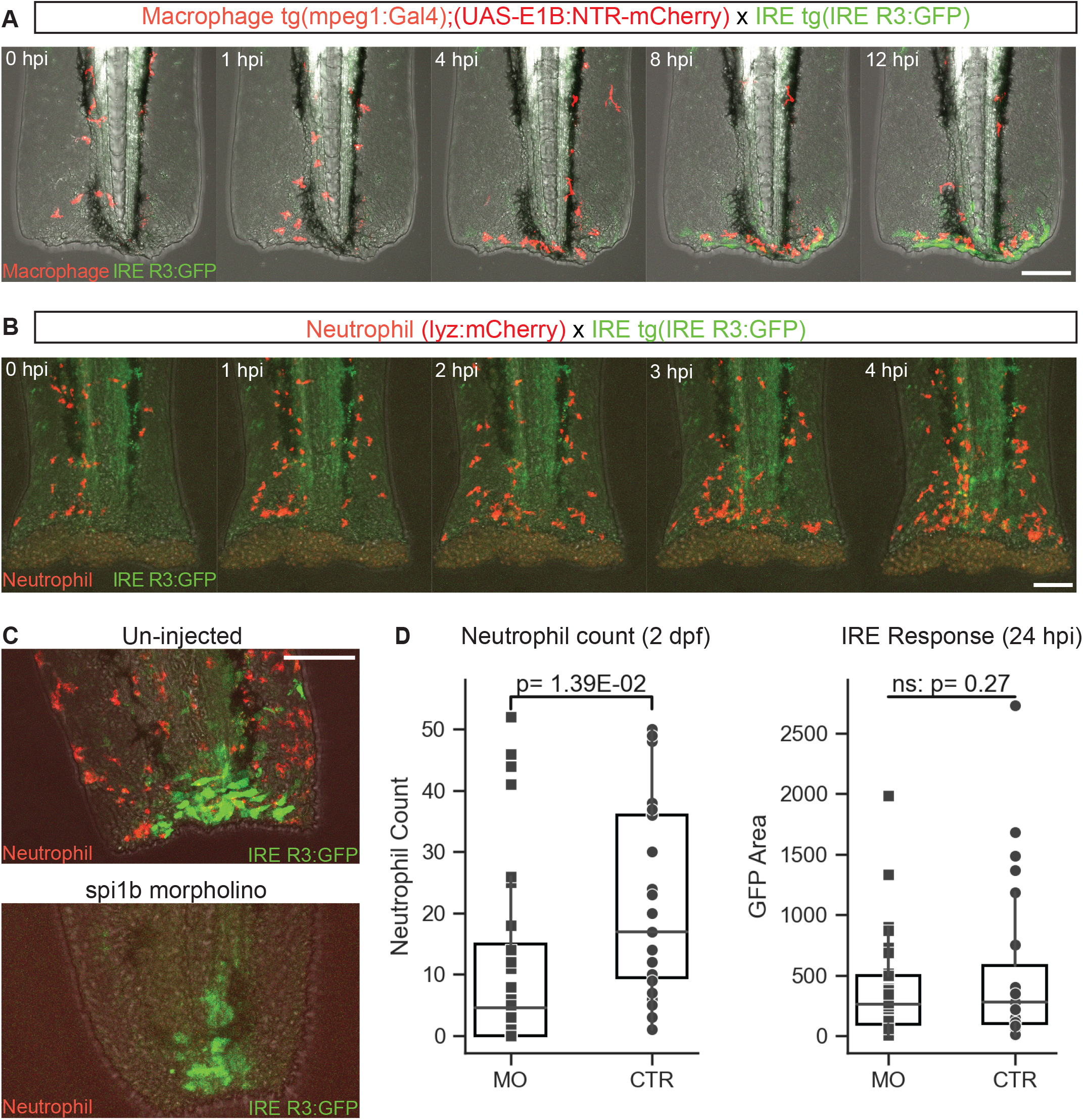
Inflammatory cells and *IRE* expression. (A) Representative images from time-lapse (Movie 1) of stable triple transgenic reporter line *tg(mpeg1:Gal4; UAS-E1B:NTR-mCherry; IRE R3:GFP)* with macrophages marked in red and *IRE R3-GFP* expressing cells in green. (B) Representative images from time-lapse (Movie 2) of double transgenic reporter line *tg(lyz:mCherry; R3:GFP)* with neutrophils marked in red and *IRE R3-GFP* expressing cells in green. Neutrophils appear in wounded tissue prior to obvious *IRE* expression while macrophages arrive concurrently. (C) Comparison of neutrophil numbers (red cells) and *IRE R3-GFP* expression (green cells) in 24 hpi, 3 dpf larvae treated with *spi1b* morpholino versus un-injected controls. (D) Quantification across multiple morpholino-treated (MO) and control (CTR) fish shows no change in the area of the *IRE R3-GFP* expression domain despite significant reduction in neutrophil count. Scale bar = 100 µm.

The larval fin fold consists of a bilayer of epithelial cells, which surround a core of mesenchymal cells and actinotrichia collagen fibers. Peripheral nerve fibers project to the fin fold area, and leukocytes enter the area following injury (Fig. 2A). Many of the *GFP*-positive cells seen after injury had the polyhedral shape typical of epithelial cells. To determine if *GFP* reporter expression was limited to the epithelium, we crossed a stable line of *tg(IRE R3:GFP)* fish to a *tg(UAS-E1B:NTR-mCherry)* line, and then crossed to *tg(ΔNp63:Gal4)* fish to mark basal keratinocytes. Dual imaging of *GFP* and *mCherry* expression showed that a subpopulation of basal keratinocytes activated the *IRE* (Fig. 2C). However, *IRE* expression was heterogenous in these cells, with adjacent cells not responding equally. In addition, *IRE* expression was clearly not limited to basal keratinocytes; with cells located beneath the epithelium also activating *GFP* expression in response to injury (Fig. 2C, arrow).

Neurons are required for regeneration after injury in many organisms including fish (Todd 1823, Goss 1956, Simões *et al*. 2014, Farkas & Monaghan 2017). To determine if neurons activate the *IRE* in response to tail injury, we imaged *tg(IRE R3:GFP; NBT:dsRed)* fish. We observed no colocalization between *IRE* expression and *dsRed* marked neural cell bodies or processes in any tails, including neurons whose cell bodies were in close proximity to other GFP+ cells (n=5) (Fig. 2D, arrow).

### 3.3 Inflammatory cells and *IRE* activation

Macrophages are also required for regeneration in many vertebrates, and are the source of numerous cytokine signals following injury (Godwin *et al*. 2013, Petrie *et al*. 2014, Wynn & Vannella 2016, Nguyen-Chi *et al*. 2017). To simultaneously visualize macrophage distribution and *IRE R3:GFP* reporter expression, tg(*IRE R3:GFP*; *UAS-E1B:NTR-mCherry*) fish were crossed to *tg(mpeg1:Gal4)* fish. Macrophages did not autonomously activate the *IRE*, as no colocalization between *GFP* and *mCherry* expression was observed (n=5 images, n=2 movies) (Fig. 3A, Movie 1). To further test if macrophages were spatially associated with *IRE* expressing cells, we used time lapse imaging to track cell movements after injury. Macrophages migrated to the site of the wound within approximately 2 hours of injury, concentrating at the wound around the same time as early *IRE* reporter expression (Fig. 3A: 4 hpi-8 hpi, Movie 1).

Neutrophils also are an early source of cytokine signaling after injury (Amulic *et al*. 2012). To compare *IRE* expression and neutrophil distribution, we crossed our GFP reporter fish to a neutrophil reporter line *tg(lyz:mCherry,cmlc2:GFP). mCherry* expressing neutrophil cells appeared in injured tails before the earliest *IRE* reporter expression (Fig. 3B: neutrophils, 1 hpi, *IRE* 4 hpi), and were less tightly localized to the site of injury compared to macrophages. Dual channel comparison showed that neutrophils themselves did not express *IRE R3:GFP* (Fig. 3B, Movie 2) (n=9 images, n=2 movies).

To test if signaling from inflammatory cells might be required for *IRE* activation, we injected fertilized eggs with a morpholino that inhibits expression of the transcription factor *spi1b*, previously shown to be required for early myeloid cell development (Rhodes *et al*. 2005). Although morpholino injected embryos showed a significant reduction in neutrophil cell numbers, this reduction did not lead to significant decrease in the area of *IRE R3:GFP* expression following injury (Fig. 3C,D). Tail resection may thus activate the *IRE* through at least some pathways that are separate or parallel to neutrophil activation, though it possible that more complete elimination of inflammatory cells would have a larger reduction in *IRE* response.

### 3.4 Identification of a minimal *IRE* element

Previous studies have shown that evolutionary conservation, chromatin accessibility, and transcription-factor binding sites may provide useful signatures to narrow the location of enhancer sequences (Mayor *et al*. 2000, Daugherty *et al*. 2017, Keilwagen *et al*. 2019). We initially hypothesized that injury responses were likely ancient, and that *IRE* activity might colocalize with the sequences in the region that are most highly conserved between mice and other organisms. Using the multi-species alignment track from the USCS genome browser, we identified 5 predominant evolutionarily-conserved subregions (designated Conserved Sequence A-E (*CSA-CSE*) (Fig. 4A). These blocks show different extents of conservation between placental mammals, monotremes, and birds. Although none of the blocks could be aligned to the zebrafish genome, we note that many enhancers show conservation of function even when the primary sequence can no longer be aligned (Fisher *et al*. 2006, Yao *et al*. 2016). We therefore concatenated the five conserved sequence blocks into a single reporter construct (*CS All*), injected it into fertilized zebrafish eggs, and assessed *GFP* expression after tail injury in the resulting mosaic transgenic larvae. Significant reporter activation was observed in 78% of larvae (Fig. 4B n= 7/9). We then built additional reporter constructs to test which of the individual sequence conservation blocks was required for *IRE* activity. Constructs missing the first, second, third, or fourth conserved regions (*CSΔA, CSΔB, CSΔC, CSΔD*) still drove robust reporter expression following injury (n= 10/15, n= 12/18, n= 9/11, and n= 10/13 transgenic larvae, respectively). In contrast, removing the fifth 1843 bp conserved region (*CSΔE)* resulted in zero transgenic larvae showing *GFP* expression following injury (Fig. 4B, n= 0/13). These results suggest that sequences located in the *CSE* region are necessary for IRE activity.

**Figure 4:**
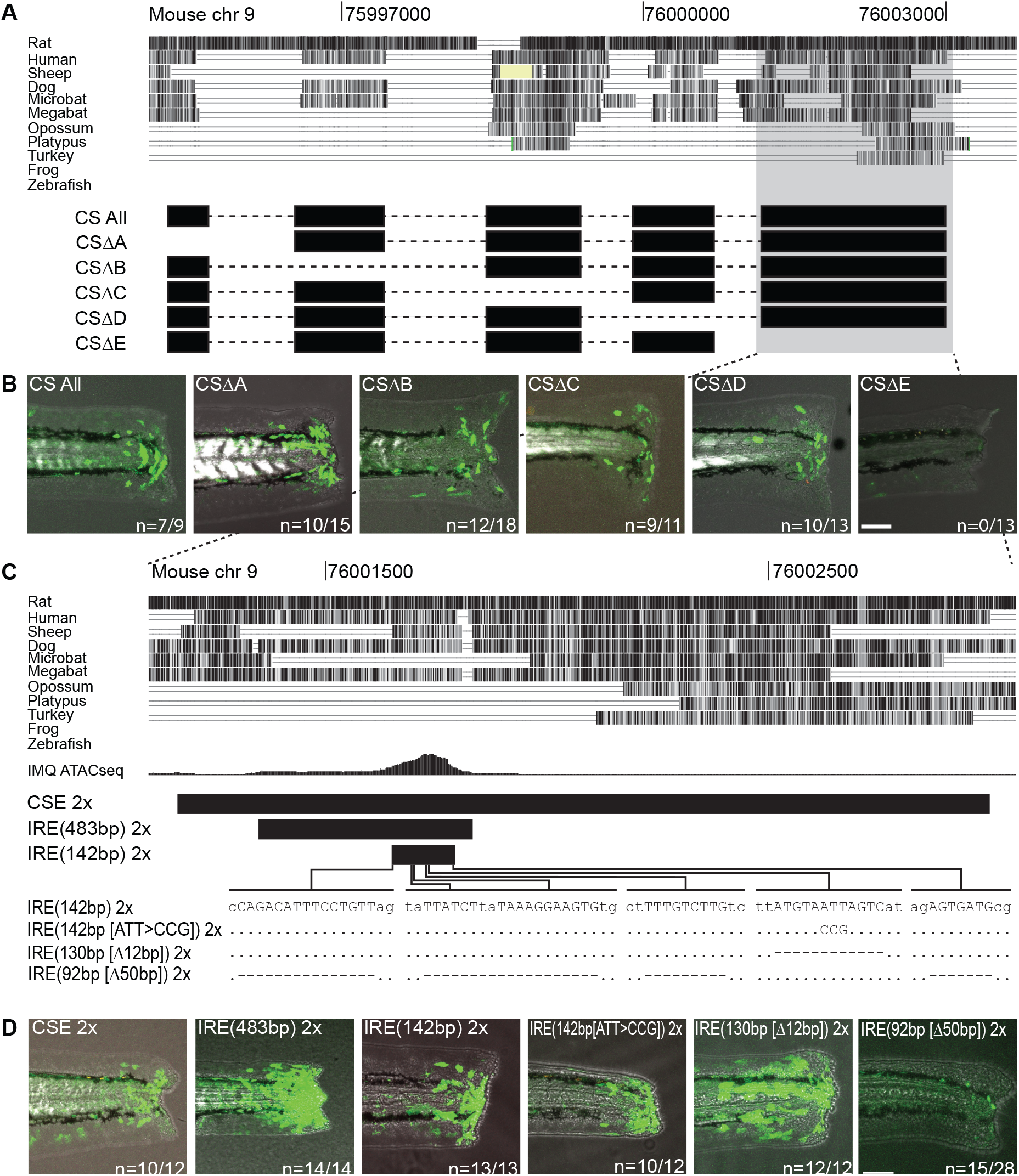
Defining a minimal injury element. (A) Schematic of the *IRE R3* sequence (mm10: chr9: 75,994,881-76,003,690). The multiple species alignment track from the UCSC Genome Browser shows 4 major regions conserved in mammals (*CSA-CSD*), and a fifth region (*CSE*) conserved between mammals and birds. Black bars show sequences used to build transgenic reporter constructs, with dashed lines indicating where the conserved regions are joined. (B) A construct containing all five conserved sequences (*CS All*) drives robust reporter expression in injured larval tails. Expression is also seen when conserved blocks A, B, C or D are deleted (*CSΔA-CSΔD*), however removing conserved sequence block E eliminates *IRE* activity (*CSΔE*). Representative expression in transient transgenic larvae is shown, with the number of fish with GFP expression indicated. (C) Detailed view of Conserved Sequence E (*CSE*) region. In addition to conservation tracks, chromatin accessible regions found in mouse epithelial cells after exposure to the proinflammatory drug IMQ are shown (IMQ ATACseq track, modified from Naik *et al*. 2017). Sequence regions used for six additional transgenic constructs are shown, the four smallest of which are identical except for the indicated base changes in the region of the IMQ ATACseq peak. (D) Robust reporter expression is seen following tail bud injury in mosaic transgenic larvae in five of the six constructs, resulting in a minimal *IRE* sequence of 130 bp. Deletion of multiple TFBS greatly reduces the extent of the injury response in the sixth construct. The number of fish with *GFP* expression indicated. All fish imaged at 4 dpf, 24 hpi. Scale bars = 100 µm.

To test if *CSE* was sufficient to activate gene expression following injury, we cloned this region individually or in tandem copies upstream of the *GFP* reporter. A single copy of *CSE* drove injury expression in 44% (n= 8/18) of transgenic larvae, and two copies drove expression in 83% (Fig. 4D n= 10/12), confirming that *CSE* has *IRE* activity.

To further narrow a minimal *IRE* region, we searched published datasets for possible changes in chromatin accessibility or transcription factor binding following injury or regeneration. Previous studies have shown that in mouse skin epithelial stem cells exposed to Imiquimod, a potent inflammatory drug, a 483 bp chromatin accessibility peak maps within *CSE* (Naik *et al*. 2017). We tested this smaller subregion of *CSE, IRE(483bp)*, and found that a 2x copy was sufficient to induce *GFP* expression following injury (Fig. 4D). By focusing on the strongest region of the chromatin accessibility peak, and by trimming sequences that appeared to be missing in the sheep genome (Fig. 4C), we further narrowed the minimal element to a 142 bp subdomain, *IRE(142bp)*, which drove robust expression in 100% (Fig. 4D n= 15/15) of fish.

Motif analysis of the *IRE(142bp)* region identified multiple predicted transcription factor binding sites (Supplemental Table S3), including a homeodomain transcription factor binding site predicted to be bound by many possible transcription factors, including some known to be expressed in the zebrafish tail after injury (Akimenko *et al*. 1995, Thummel *et al*. 2007). To test if this binding site was necessary for reporter function, we altered base pairs within the predicted binding motif (*IRE(142bp [ATT>CCG])*, or deleted the site entirely *IRE(130bp [*Δ*12bp])*. Robust expression was visible following injury with both constructs (Fig. 4D n = 10/12 and n=12/12, respectively). Thus, the predicted homeodomain transcription factor binding site is not required for *IRE* activity, and a 130 bp sequence is sufficient to drive reporter expression after tail injury.

To test the potential role of the other predicted transcription factor binding sites, we deleted the remaining five sites in a single construct (*IRE*(92bp [Δ50bp]). When compared to *IRE(142bp*) the extent of expression was markedly reduced, though some green cells were still detected in a fraction 54% (n= 15/28) of fish (Fig. 4D). This suggests that the targeted transcription factor binding sites are required for full *IRE* activity, but that other sequences within the *IRE* may also contribute.

### 3.5 The minimal *IRE* drives expression in multiple cell types and injury models

Previous studies have shown that large injury response regulatory regions may consist of either single elements that respond in multiple tissues, or clustered but distinct enhancers that each respond to injury in particular tissues (Yang & Kang 2019). We therefore compared the ability of the large and minimal *IRE* region to drive expression in two different injury models.

In the larval fin, both large and small *IRE* constructs drove reporter expression in clipped tails, including in both the epithelium and mesenchymal core of the fin (Fig. 5 C-D, large *IRE R3*, n= 4/4; *IRE(142bp)*, n= 5/5; and *IRE(130bp [*Δ*12bp]*, n= 4/4).

**Figure 5:**
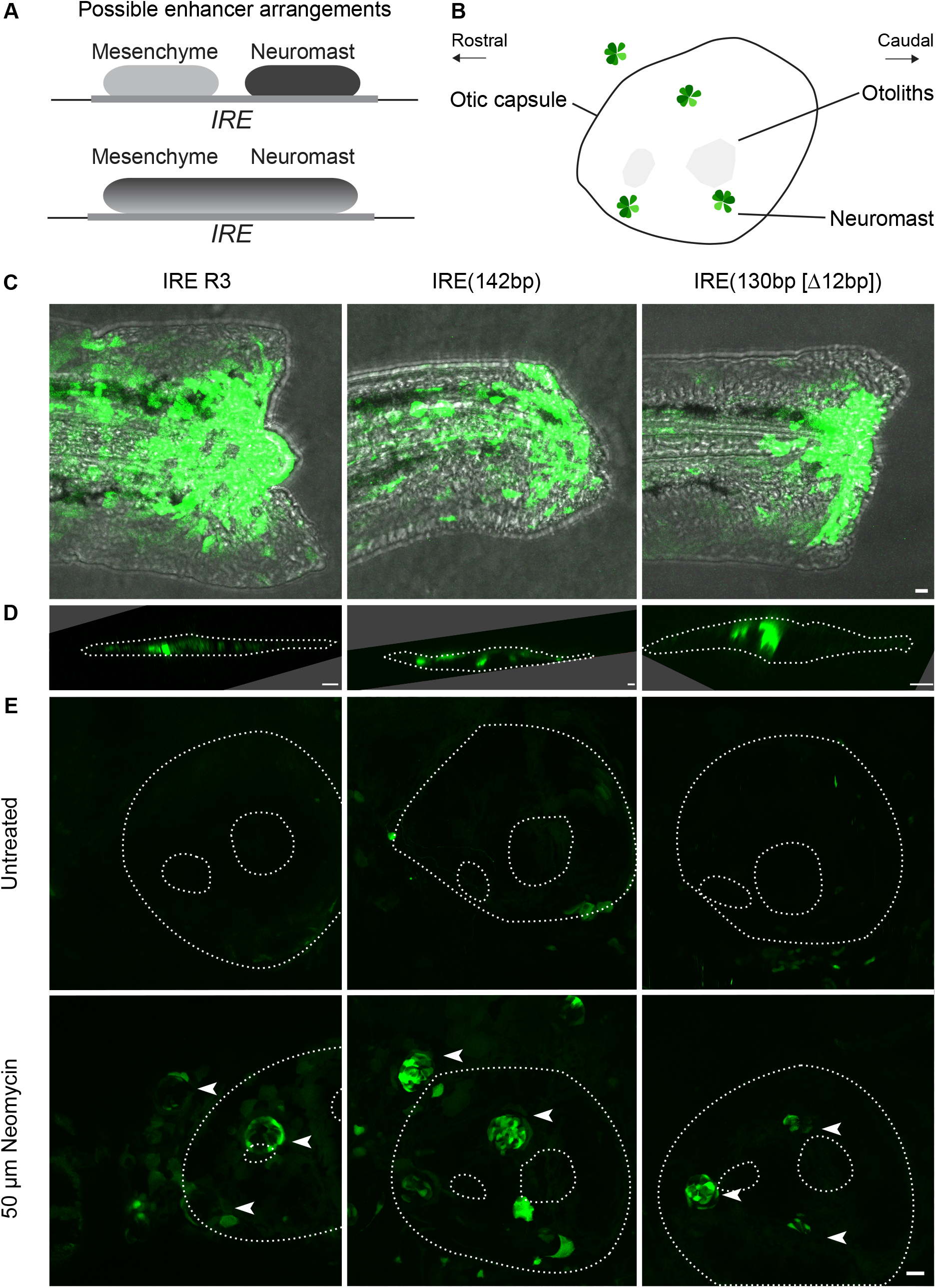
A minimal *IRE* responds in both mesenchymal and epithelial tissues. (A) Different models of arrangements are possible in injury control regions, including separate response elements for different tissues (top), or a single response element for multiple tissues (bottom). (B) Schematic of the stereotyped neuromast locations found superficial to the zebrafish otic vesicle (circle) and embedded otoliths (grey). The otic vesicle serves as a convenient landmark to identify these neuromasts. (C,D) Both large and small *Bmp5 IRE* constructs (*IRE R3, IRE(142bp), IRE(130bp [Δ12bp])* drive expression in injured tail fins, including mesenchymal cells seen in the core of the fin (Panel D). (E) The same constructs, including the smallest, also drive *GFP* reporter activity in zebrafish neuromasts (arrowheads) following injury with 50 µM neomycin exposure, but not in untreated 6 dpf fish. Scale bars: C, D, E = 20 µm.

To test responses to chemical injury rather than tail resection, we also examined *IRE* responses after lateral line injury. The lateral line of zebrafish is a mechanosensory organ used by fish to sense water movements (Mogdans 2019). The sensory apparatus is made up of neuromasts, cone shaped structures dispersed along the lateral flank of the fish that contain innervated sensory hair cells surrounded by rings of support cells and mantle cells (Webb 2011). Just as hair cells in the mammalian inner ear are known to be sensitive to killing by aminoglycoside antibiotics like neomycin and gentamycin, lateral line hair cells in fish can also be killed by the same compounds that are ototoxic in humans (Harris *et al*. 2003). To determine if the *IRE* is activated after hair cell injury, we incubated developing transgenic larvae with neomycin. Reporter expression was observed in the support, mantle, or remaining hair cells of the neuromast following neomycin treatment of transgenic embryos. Expression was observed in fish carrying either the larger *IRE R3* construct, or the much smaller *IRE(142bp)* and *IRE(130bp [Δ12bp])* constructs compared to control (Fig. 5E, n= 18/18; n= 20/24; n= 12/15). These results indicate that the same minimal *IRE* sequence is capable of driving expression in multiple cell types and injury models.

## 4 Discussion

Many key signaling molecules used to control tissue formation during embryonic development are also expressed at sites of tissue injury. Previous studies showed that re-expression of the *Bmp5* gene after injury is controlled by a regulatory region distinct from the developmental enhancers that program highly patterned expression during embryogenesis (Guenther *et al*. 2015). Although *Bmp5* has become a canonical example of injury control by a dedicated regulatory region (Rodriguez & Kang 2019, Yang & Kang 2019), the time and expense of generating and scoring transgenic mice made it difficult to further narrow the large injury response region using previous approaches.

Here we show that the mouse *IRE* region is also activated in a variety of zebrafish injury models. These results indicate that the *IRE* responds to ancient injury sensing mechanisms that are shared in organisms diverged by hundreds of millions of years. Interestingly, conservation of *IRE* functional response occurs even though the primary sequence of the mouse *IRE* can no longer be directly aligned to the zebrafish genome. Similar results have been reported for many other enhancers, such as the human *RET* enhancer which functions in zebrafish, or the mouse and acorn worm *SBE1* enhancers which also retain function in the absence of primary sequence alignment (Fisher *et al*. 2006, Yao *et al*. 2016). In each of these cases, enhancer function is likely preserved due to similar presence of the necessary transcription factor binding site (TFBS) motifs, despite potential re-arrangements and other changes in sequence. This is consistent with conserved expression patterns but rapid turnover of transcription factor binding sites that have also been observed between even closely related drosophila species that diverged on time scales much shorter than the millions of years separating mouse and zebrafish (Ludwig *et al*. 1998, Moses *et al*. 2006, Swanson *et al*. 2011).

Many features of the zebrafish response recapitulate key *IRE* properties known in mice. For example, the original *IRE* drove expression in multiple injury models and multiple cell types in mouse experiments, including mesenchymal cells around fractured bones, lung epithelial cells following chemical injury, and basal epithelial cells following skin punch biopsy (Guenther *et al*. 2015). Similarly, we find that the *IRE* region can also be activated by both mechanical and chemical injuries in zebrafish, and is induced in multiple cell types, including tail fin epithelium and mesenchyme, and in mechano-sensory and support cells in neuromasts. *IRE* response has not previously been characterized in mouse mechanosensory cells after injury. However, we note that the lateral line of fish and the inner ear of mammals are made of homologous cell types that share sensitivity to aminoglycoside antibiotics. Mouse *Bmp5* is known to be upregulated in regenerating sensory epithelia of the inner ear after gentamycin-induced injury (Bai *et al*. 2019). Similarly, zebrafish *bmp5* is expressed in the stem cell niche responsible for maintaining neuromast cells (Lush *et al*. 2019) and expression of *bmp5* has previously been shown to be induced after hair cell injury by neomycin (Jiang *et al*. 2014). Our studies add to the growing evidence that the *IRE* can activate gene expression in multiple contexts following diverse types of injury.

Other enhancers that are active in multiple injury models have subsequently been shown to consist of separate enhancers in close physical proximity, each activating expression in different locations (Kang *et al*. 2016). The original mouse *IRE* was mapped to a large 17.8 kb region, raising the strong possibility that its diverse activities might also be explained by multiple linked enhancers, perhaps one activated by bone fracture, and others by lung injury, skin injury, etc. Using our ability to rapidly screen constructs in the zebrafish injury models, we have now been able to narrow the physical size of the *Bmp5* injury response region by more than a hundred-fold. Importantly, even a minimal 142 bp *IRE* sequence can still be activated by both mechanical and chemical injuries, and still drives induced expression in multiple cell types, including tail epithelium, tail mesenchyme, and neuromasts. Although it is possible that even this small region consists of multiple different enhancers, we favor the hypothesis that *IRE* activity is due to a single functional element that responds to a damage signal shared by many types of injury.

What might this shared signal be? Previous studies of hair cell injury show that aminoglycosides enter hair cells through mechano-electric transduction channels and activate cell death signaling. The precise mechanisms inducing cell death are not completely known, but in fish, caspase-independent apoptosis is known to play an important role (Coffin *et al*. 2013). This apoptosis is likely triggered by loss of membrane integrity of the mitochondria and concomitant increase in reactive oxygen species, although other mechanisms may also contribute (Karasawa & Steyger 2011, Coffin *et al*. 2013, Esterberg *et al*. 2016). Reactive oxygen species are also known to be induced by mechanical injury like tissue resection. For example, a gradient of hydrogen peroxide is rapidly generated by DUOX activation after tail clipping in zebrafish, and this gradient has been implicated in the attraction of leukocytes to the wound site (Niethammer *et al*. 2009). Our time lapse movies show that *IRE* driven *GFP* expression is first apparent near the wound edge within the first hours after injury, with gene expression then increasing and spreading across several cell diameters. Although immune cells are clearly activated at larger distances, *IRE* is not induced within these distant cells, or in distant epithelial and mesenchymal neighbors. *IRE* activation thus most likely occurs through a rapidly acting, local signal triggered by diverse types of injury. In the future it would be interesting to simultaneously visualize the pattern of *IRE* activation with a variety of candidate upstream signals, including calcium flux, reactive oxygen species, tissue tension detected by adhesion molecules, osmotic shock, or changes in transepithelial electrical potential (Rieger & Sagasti 2011, Yoo *et al*. 2012, Gault *et al*. 2014, Enyedi *et al*. 2016, Kennard & Theriot 2020).

The small 142 bp sequence that we have now identified with *IRE* activity should also aid the study of molecular mechanisms leading to injury-induced activation. The minimal *IRE* sequence contains predicted binding sites for multiple classes of upstream factors with plausible connections to tissue repair and regeneration. Comparison of *IRE*-*142bp* with the UNIPROBE database (Newburger & Bulyk 2009) of experimentally measured binding interactions identifies predicted binding motifs for homeodomain transcription factors, Elf3 and Elf5, Gata3 and Gata6, Sfpi1, Sox14, Smad3, and Pbx1 (Supplemental Table S3). Elf3 is an epithelial-specific trans-activator that is known to be expressed following H_2_O_2_ treatment in zebrafish (Lisse *et al*. 2016). *Gata3* is induced after traumatic brain injury in adult zebrafish, is required for caudal fin regeneration (Kizil *et al*. 2012), and has been shown to regulate *Bmp5* expression in prostate epithelial stem cells (Tremblay *et al*. 2020). *Gata4* expression is important in zebrafish heart regeneration (Kikuchi *et al*. 2010). Additionally, multiple *Sox* genes, including *Sox14* are known to be induced in zebrafish spinal cord injury (Ghosh & Hui 2016). *Pbx1* has a less clear role in injury response, but is slightly downregulated in caudal fins after injury (Mathew *et al*. 2009). Smad3 binding sites are known to be important in the *careg* regeneration associated enhancer element (Pfefferli & Jaźwińska 2017) in zebrafish. Their presence in the *IRE* may suggest a common gene network with *careg* as well as raising the intriguing possibility that *Bmp5* expression is autoregulatory.

We tested the effect of removing the single binding site with the largest number of candidate upstream factors in the current study. Complete deletion of the predicted binding site for numerous homeodomain transcription factors produced a smaller *IRE* 130 bp sequence that could still be activated in both tail fin and neuromast injuries. Extending this approach to the other classes of predicted binding sites suggests that multiple transcription factors may act in concert to produce *IRE* activation. However, further investigation of these sites individually and in combination is needed to further characterize the specific roles of each binding site in multiple wound models.

Rapid expansion of comparative sequencing approaches in mammals may help prioritize particular subregions of the *IRE* for additional studies. For example, the Zoonomia project has recently shown that genome data from 240 different mammals can help identify particular base pairs most likely to be important in predicted binding sites for upstream factors (Zoonomia Consortium 2020). Comparison of the 142 bp *IRE* region across diverse mammals reveals several interesting patterns (Supplemental Figure S3). First, although the *IRE* sequence is generally well conserved in mammals, several species have lost portions of the proximal sequence. When designing constructs, we used partial loss of the *IRE* region in the sheep genome to help refine the proximal side of the minimal *IRE* region. Similar reasoning with additional genome sequences suggest that further narrowing of the *IRE* region should be possible. The best conserved binding site in the entire 142 bp *IRE* is the predicted Gata3/Gata4 response region, which is also the only portion of the *IRE* region still present in two sequenced sloth species. Although the status of injury repair in sloths is poorly characterized, the overall conservation pattern of the *IRE* in mammals, together with recent studies showing that GATA3 can act as a direct regulator of *Bmp5* expression in prostate stem cells (Tremblay *et al*. 2020), makes GATA a particular promising candidate for an upstream regulator of the *Bmp5* injury response.

Interestingly, the mammalian comparisons suggest that *Bmp5 IRE* sequence is entirely missing in multiple microbat species. The absence of the sequence is not due to genome assembly errors, as PCR amplification and sequencing across the region confirms a ∼500 bp deletion that completely removes the *IRE* region (Methods). In contrast, the *IRE* is still retained in megabats, a sister group of flying mammals with larger body sizes in the Chiroptera group. Interestingly, microbats are unusually long-lived relative to their size (Wilkinson & Adams 2019). Several metabolic and genomic studies have suggested that microbats may have evolved unique mechanisms to reduce production of reactive oxygen species, likely as a physiological adaption to tolerate the high metabolic demands of powered flight (Brunet-Rossinni and Austad 2004; Z. Huang, Jebb, and Teeling 2016; Kacprzyk et al. 2017, but see Wilkinson and Adams, 2019). Bats are also unusual among mammals in completely losing multiple *PHYIN* genes, which activate inflammasome and interferon pathways in response to free DNA; the *IL36G* gene, a pro-inflammatory interleukin; and the *LRRC70* gene, which amplifies cellular responses to bacterial lipopolysaccharide and multiple cytokines (Ahn *et al*. 2016, Jebb *et al*. 2020). Loss of the *IRE* sequence may represent another example of a genomic loss in microbats which helps prevents hyperactivation of inflammatory and repair pathways in an animal group with unusually large metabolic activity.

### Summary

The mechanisms that lead to activation of BMP expression at fracture sites and other types of tissue injury are still poorly understood. Our studies have greatly narrowed the injury response control sequences in the *Bmp5* gene. The same *IRE* region is induced by diverse types of injury in organisms as distinct as zebrafish and mice, indicating that a single small injury enhancer can respond to ancient upstream mechanisms that act locally near many types of tissue damage. Further study of the candidate pathways identified in this study should help identify specific molecular regulators of the *IRE* and may suggest new mechanisms for modulating BMP expression in both bone fractures and many other types injury.

## 5 Acknowledgments

We thank Tuky Reyes, Chenelle Hill, and Virginia Herbert for fish husbandry; Karen Mruk and Andrew Kennard for providing fish stocks; Samantha Larsen for providing IMQ peaks; Heidi Chen, Karthik Jagadeesh, and Gill Bejerano for assistance calling transcription factor binding sites; a Mount Desert Island Biological Laboratory course for useful training in injury models; Catriona Logan, Kara Bower, Kayla Huemer, and Emma Li for useful discussions; Harini Iyer, and Dan Lysko, Paola Sidoli; Garrett Roberts-Kingman, Janet Song, and all members of the Talbot and Kingsley laboratories for reagents, technical assistance with fish procedures, and helpful discussions and comments on the manuscript. This work was supported in part by training and research grants from the National Institute of Health (T32 predoctoral program 2T32GM00779038 (ISH), NIH R35 NS111584 (WST) and AR42236 (DMK)). DMK is an investigator of the Howard Hughes Medical Institute.

## Notes

### Competing Interest Statement

The authors have declared no competing interest.

