## Supplementary Materials for "Characterization of mouse *Bmp5* regulatory injury element in zebrafish wound models"

**Supplementary figures included in this file:**
Supplemental Figure 1. *Bmp5* injury reporter expression in adult fin ray breaks.
Supplemental Figure 2. Empty expression vector control.
Supplemental Figure 3. Alignment of the minimal 142 bp IRE sequence across 242 mammals sequenced by the Zoonomia Consortium
(Owing to its large size, Fig. S3 can be accessed for larger viewing here)

**Supplemental tables 1-4 provided in separate Microsoft Excel file (*also available here*):**
Supplemental Table 1: Primers and oligos used in this study
Supplemental Table 2: Regions covered by different *Bmp5 IRE* constructs
Supplemental Table 3: UNIPROBE analysis of transcription factor binding sites in IRE142
Supplemental Table 4: UNIPROBE analysis of engineered mutations in IRE(142bp)

**Supplemental movies provided in separate avi files:**
Movie 1: Macrophages and IRE expression. *also available* here
Movie 2: Neutrophils and IRE expression. *also available* here

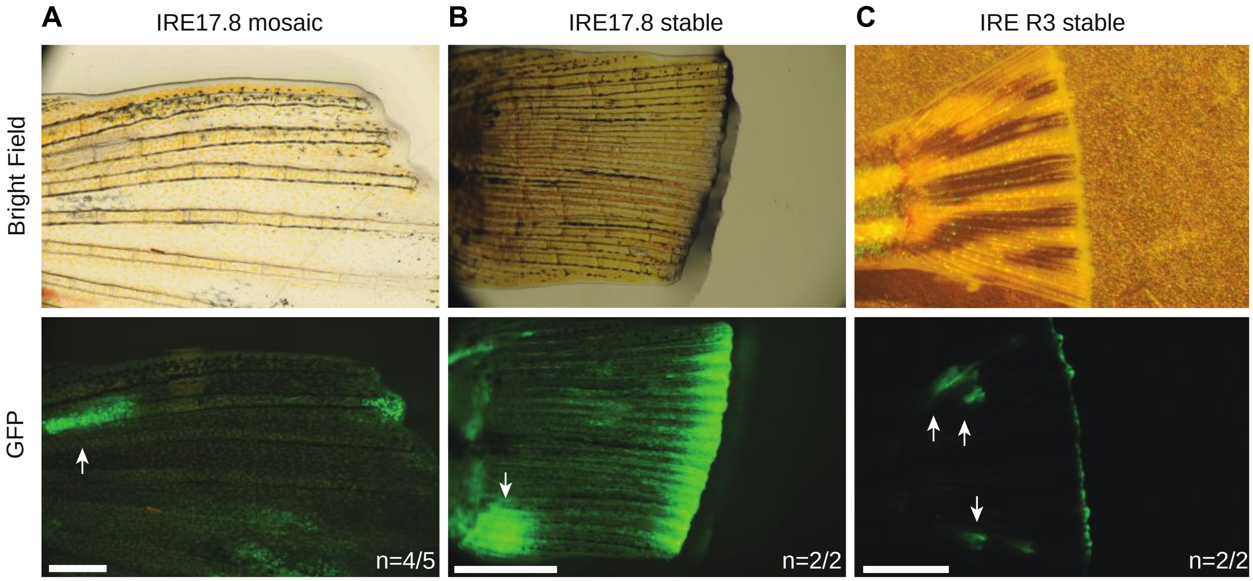


Supplemental Figure 1. *Bmp5* injury reporter expression in adult fin ray breaks.

(A) Injured adult caudal fin in mosaic *tg(IRE17.8:GFP)* fish showing the location of injury and *GFP* expression at 72-hours post injury. (B) Additional image of stably transgenic *tg(IRE17.8:GFP)* fish showing GFP reporter expression at the wound edge at 72-hours post injury. (C) Injured adult caudal fin in stable line of *tg(IRE R3:GFP)* fish at 72-hours post injury. (A-C) Incidental breaks in the hemiray bones also show GFP expression (white arrows). Number of fish with *GFP* expression indicated. Scale bars: A =1 mm, B, C = 3 mm.

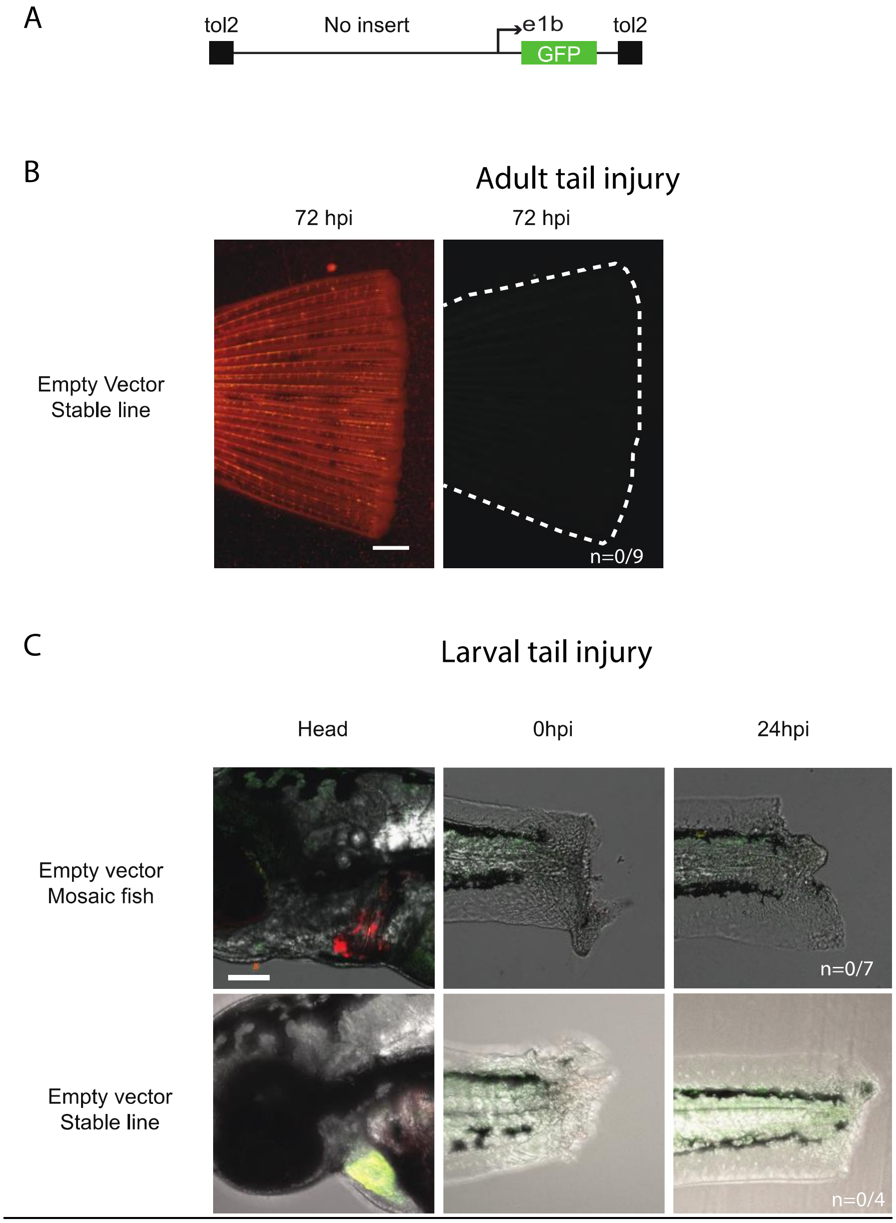


Supplemental Figure 2. Empty expression vector control.

(A) Schematic of the zebrafish expression vector. (B) Representative caudal fin showing the location of injury and absence of GFP expression from the empty vector at 72 hours-post-injury in the adult (C) Representative larval zebrafish. Transgenic mosaic fish are screened for *cmlc2:mCherry* expression (red) in the heart comparable to *cmlc2:mCherry* expression in a stable line (yellow). The empty vector did not drive expression in injured tails N= GFP positive fish/ total stable (B, C *bottom*) or mosaic (C *top*) transgenic fish screened. Scale bars: B= 5 mm, C=100 µm.

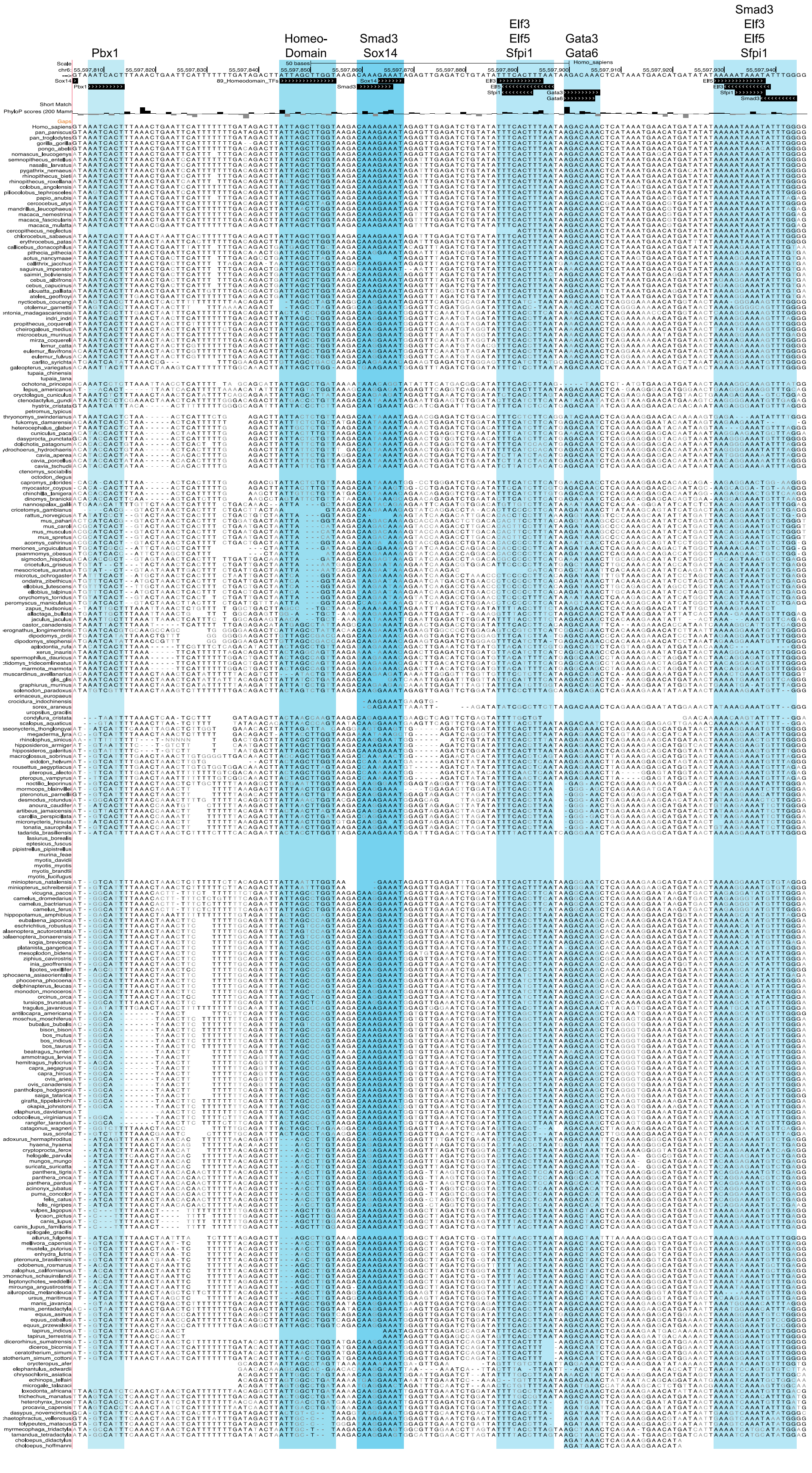
Pbx1
Homeo-Domain
Smad3 Sox14
Elf3 Elf5 Sfpi1
Gata3 Gata6
Smad3 Elf3 Elf5 Sfpi1

Supplemental Figure 3. Alignment of the minimal 142 bp *IRE* sequence across 242 mammals sequenced by the Zoonomia Consortium.

PhyloP conservation scores are shown at individual base pair resolution. The locations of six major classes of predicted transcription factor binding sites discussed in the text are shaded in blue. A 12-bp engineered deletion removing the homeodomain binding site does not block *IRE* activation after injury (see Results). Eight different microbat species are missing the entire *IRE* region (*Lasiurus borealis*, *Eptesicus fuscus*, *Pipistrellus pipistrellus*, *Murina feae*, *Myotis davidii*, *Myotis myotis*, *Myotis brandtii*, and *Myotis lucifugus*), while two sloth species (C*holoepus hoffmanni,* C*holoepus hoffmanni)* retain only the region that includes the predicted binding site for GATA family.
